# Bambara Groundnut Rhizobacteria Antimicrobial and Biofertilization Potential

**DOI:** 10.1101/2020.02.27.964346

**Authors:** Caroline F. Ajilogba, Olubukola O. Babalola, Patrick Adebola, Rasheed Adeleke

## Abstract

Bambara groundnut, an underutilized crop has been proved to be an indigenous crop in Africa with the potential for food security. The rhizosphere of Bambara groundnut like other legumes contains several important bacteria that have not been explored for their plant growth-promoting properties. The aim of this research was to determine the potentials of rhizobacteria from Bambara groundnut soil samples as either biofertilizer or biocontrol agents or both to help provide sustainable agriculture in Africa and globally. Analyses of Bambara groundnut rhizospheric soil samples included chemical analysis such as nitrogen content analysis using extractable inorganic nitrogen method as well as cation exchangeable capacity using ammonium acetate method. Plant growth-promoting properties of isolated rhizobacteria tested include indole acetic acid, hydrogen cyanide, phosphate solubilization, 1-aminocyclopropane-1-carboxylate and ammonia production activities using standard methods. In addition, antifungal assay dual culture method was used to analyze the biocontrol properties of the isolates. Phylogenetic analysis using 16S rRNA was also carried out on the isolates. Isolated rhizobacteria from bambara groundnut rhizosphere were cultured. All the isolates were able to produce ammonia and 1-aminocyclopropane-1-carboxylate while 4.65%, 12.28% and 27.91% produced Hydrogen cyanide, Indole acetic acid and solubilized phosphate respectively, making them important targets as biocontrol and biofertilizer agents. The growth of *Fusarium graminearum* was suppressed *in vitro* by 6.98% of the isolates. Plant growth promoting activities of rhizobacteria from bambara groundnut rhizosphere reveals that it has great potentials in food security as biofertilizer and biocontrol agent against fungal and bacterial pathogens.

## 1.0 Introduction

The use of chemicals to inhibit the growth of pathogenic microorganisms in plant disease control has been a global issue. Research for more healthy environmental control methods have led to biocontrol and biofertilization. Rhizospheric soils of legume crops have been considered a reservoir for plant growth promoting rhizobacteria (PGPR). Bambara groundnut (*Vigna subterranean* L. Verdc), a legume crop, is one of the neglected and underutilized (NUS) species. The term ‘NUS’ is used to mean wild species of plant which are non-commodity cultivated. They form part of a large agro biodiversity portfolio that are not used as a result of an array of factors such as agronomic, genetic, economic, social and cultural factors [1]. They are traditionally grown by subsistence farmers in their various localities where they are useful in supporting and securing nutrition in local communities in order to meet their socio-cultural needs and traditional uses. They have been largely ignored by research and development and so there is no competition for them compared to other well-established major crops. This results in the loss of both their diversity and traditional knowledge. It is a food known as a balanced diet as it contains the right proportion of protein (16.25%), carbohydrate (63%) and fats (6.3%) [2]. The protein is high in both lysine (6.6%) and methionine (1.3%) [3]. Because of its richness in protein and the fact that it is nutritious, it is a source of food security especially for small scale farmers and small households [4]. Bambara groundnut is also very rich in micronutrients such as potassium, calcium and iron with a high proportion of fiber [5]. There are different varieties with varying mineral composition for example the red varieties contains iron twice as much as the cream variety making it quite suitable for mineral deficient in iron [4]. It was observed that fermentation of bambara groundnut helped to improve its mineral composition which invariably reduced the different factors that inhibited nutrient utilization such as trypsin, oxalate, phytic and tannic acid [6].

Bambara groundnut has the ability to grow under different climatic and soil conditions that are harsh and extreme thus making it suitable to be grown in semi-arid lands. The soil rhizosphere of legumes which include bambara groundnut has been indicated to enhance plant growth and also for controlling plant pests and diseases. Such beneficial attributes are associated with a host of rhizobacteria that inhabit this rhizosphere and are also sometimes referred to as PGPR. Examples of such bacteria include *Bacillus* spp, *Actinomycetes*, spp *Pseudomonas* spp, *Burkholderia* spp and *Rhizobium* spp [7]. The diversity of these microbial communities is driven by plant-microbe activities such as organic compound secreted by plants as well as availability and quantity of nutrients released by microbes. These interactions also play a crucial influence on the health, growth, pest and disease susceptibility of the plant as well as the health of the soil [7].

Plant growth-promoting rhizobacteria (PGPR) also promote plant growth directly by producing phytohormones such as IAA and HCN. Sometimes they enhance iron chelation (siderophore production) and supply of nutrients such as phosphorus (phosphate solubilization) and nitrogen (nitrogen fixation) to also promote plant growth. They are well known to participate in biofertilization, which involves enriching rhizospheric soil, making nutrients available to the plants as well as aiding the plants in nutrient uptake and the subsequent use of the nutrients for metabolic processes by the plants [8]. They have also been found to help in biocontrol of plant pests and diseases by suppressing and/or inhibiting the growth of pathogens in/on plants [9]. As new pathogens causing plant diseases are being identified, there is the need to find better bioalternatives that have not been harnessed but have prospects. The rhizosphere of bambara groundnut has not been explored like other legumes for rhizobacteria that are important in biofertilization and biocontrol. This study aimed at evaluating the rhizobacteria found in the rhizosphere of Bambara groundnut for their biofertilization and biocontrol potentials as a tool for food security.

## 2.0 Materials and methods

### 2.1 Planting of Bambara groundnut

The propagation of Bambara groundnut was through seeds on level seedbed or ridges where the soil is wet. Seeds from two different landraces VL and VBR were planted on seedbeds in plots that are 50 cm apart and spacing between seed holes on each plot was 50 cm apart (Fig. 1a and Fig 1b). Seeds were planted 3-4 cm deep in the soil. Twenty-five (25) plots were cultivated for each replication and the experiment was repeated thrice.

**Figure 1:**
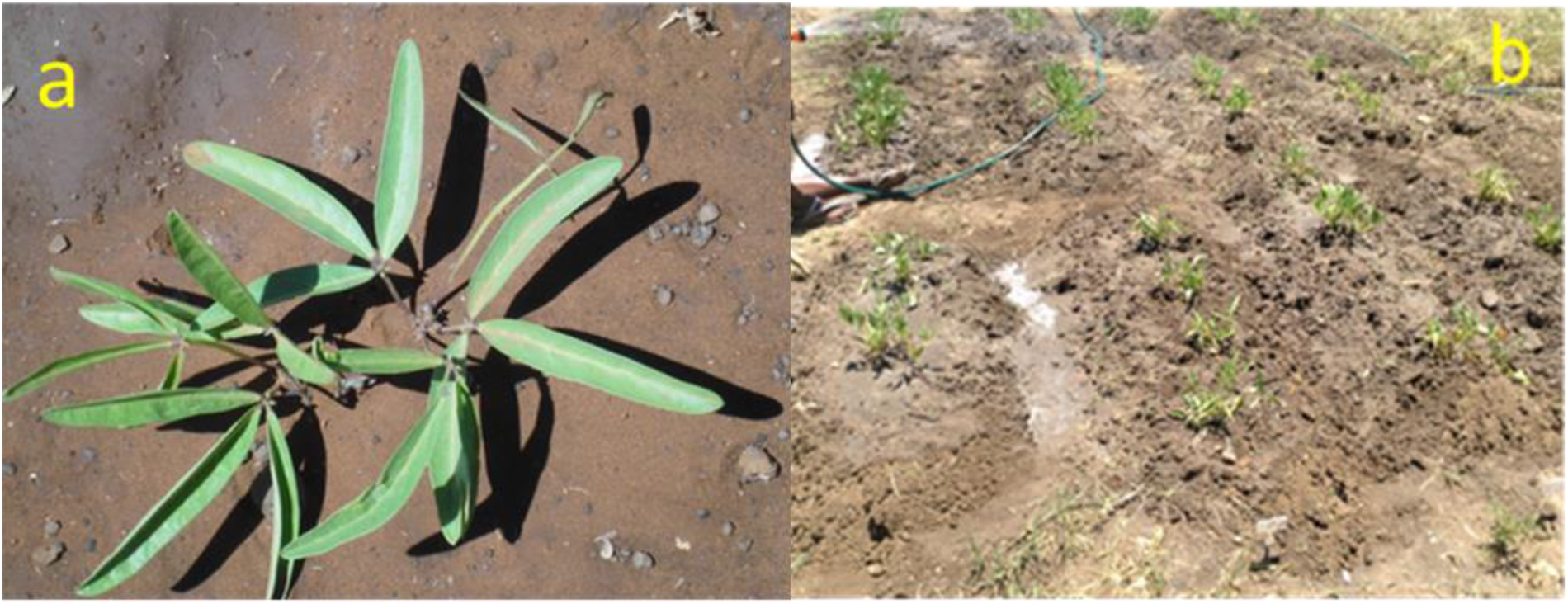
Plot showing Bambara groundnut 2 weeks after planting (a and b)

### 2.2 Soil sampling and collection

Soil samples were collected from field trials during the planting period between October 2014 and March 2016 from the North-West University Agricultural Farm, Mafikeng Campus (Lat. 25°78’91 “ Long. 25°61’84) Mafikeng, South Africa. Four soil samples were collected randomly from the uprooted Bambara groundnut root rhizosphere at four weeks interval from bulk soil before planting for 16 weeks corresponding to different growth stages of the plant. Twenty four (24) soil samples were collected in all to a depth of 15 cm per sample. Samples were collected in triplicates from the rhizosphere of two different landraces and were stored at 4°C until ready for use.

### 2.3 Soil Analysis

According to the International Standard Organization (ISO) standard 11464, the samples were prepared for analysis by drying at room temperature, pulverized, and sieved through a 2 mm sieve. All glassware used for soil analyzes were washed thoroughly, soaked in 20% nitric acid and rinsed with deionized water to prevent the presence of impurities. The selected physical and chemical parameters of the samples were analyzed using standard laboratory procedures [10].

All soil analysis were repeated twice.

#### 2.3.1 Physical and chemical analysis of samples

##### 2.3.1.1 Determination of CEC and extractable cations

Exchangeable cation was determined using the ammonium acetate method [11]. Sample of the soil was aliquot and sieved using a 2-mm screen and allowed to air dry (at a temperature of ≤60°C). Ten (10) g of the air-dried soil was placed in a 500-mL Erlenmeyer flask and 250 mL of neutral, 1 N NH4OAc was added to it. The flask was thoroughly shaken and allowed to stand overnight. The soil was filtered with light suction using a 55-mm Buchner funnel. The soil was leached with the neutral NH4OAc reagent until no test for calcium can be obtained in the effluent solution. The soil was then leached four times with neutral 1 N NH4Cl and once with 0.25 N NH4Cl.

The electrolytes were washed with 99% isopropyl alcohol having a volume of between 150 to 200 mL. When the test for chloride in the leachate (use 0.10 AgNO_3_) became negligible, the soil was allowed to drain thoroughly. The adsorbed NH4 was determined by the aeration method [12].

##### 2.3.1.2 Nitrogen analysis of soil samples

Soil samples were dried at 80°C, ground to a powder and 1 g analysed for nitrogen (N) by Kjeldahl digestion [13].

##### 2.3.1.3 Determination of nitrate composition

Nitrate contents of the samples were determined using the equilibrium extraction method. Ten (10) g ≤ 2.0 mm of air-dry soil was placed into a 250 mL wide mouth extraction bottle and 100 mL of 0.1 mol dm^-3^ of KCl was added, stoppered and shaken for 30 min on a shaker. The solution obtained was filtered to get a clear extract and the nitrate contents were determined in the clear extract.

##### 2.3.1.4 Determination of pH and redox potential

Ten (10) g of soil was weighed and mixed with 25mL of distilled water to obtain ratio 1:2.5 (m:v) soil-water suspension and left to shake for 1 h and left standing overnight for pH measurement with a pH meter Jenway 3520^™^ (Lasec, South Africa). The pH meter was standardized using calibration buffer 4, 7 and 9. The combined electrode was inserted into supernatant and pH values and redox potential of the samples was recorded, electrode was washed with distilled water after each reading.

##### 2.3.1.5 Determination of organic matter

Organic carbon was determined by Walkley-Black method [14]. One (1) g of air-dried sample was weighed and 10 mL of 1 N potassium dichromate solution was added followed by addition of 20 mL concentrated sulphuric acid and the beaker was swirled to mix the suspension. The solution was left undisturbed and allowed to stand for 30 min and 200 mL deionized water and 10 mL concentrated orthophosphoric acid was added. Twelve (12) drops of diphenylamine indicator (1 g diphenylamine in 100 mL concentrated sulfuric acid) was added with continuous stirring on a magnetic stirrer and finally the mixture was titrated with 0.5 M ferrous ammonium sulphate until a colour change from violet-blue to green was observed. A blank solution was also prepared in which no sample was added. The percentage of organic matter was used to calculate the organic carbon content by using the conversion factor 1.724 and the fact that 58% of the soil organic matter is the average content of carbon [14]. This calculation is given below;

Percentage organic matter in soil is calculated as written below:

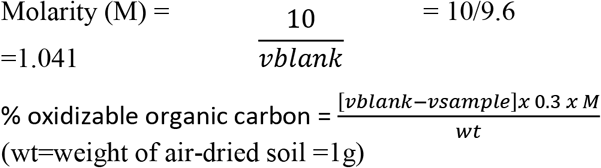

% total organic Carbon (w/w) = 1.334 x % oxidizable organic carbon

% organic matter (w/w) =1.724 x % total organic Carbon

### 2.4 Preparation of soil samples for bacterial isolation

Soil samples from the rhizosphere of Bambara groundnut were collected. Samples were prepared according to Abdulkadir and Waliyu [15] and placed into sterile 50-mL Falcon tubes (Becton Dickson Paramus, N.J) and kept on ice or at 4°C until it was needed (within 3 days).

### 2.5 Culturing and isolation of bacteria from soil samples

Isolation and enumeration of bacteria present in the soil sample were performed by serial dilution plate technique using tryptone soy agar (TSA) and the procedure of Cavaglieri, Orlando, Rodriquez, Chulze and Etcheverry [16]. A 10-fold dilution series was prepared from the rhizospheric soil of bambara groundnut in sterile distilled water and 0.5 mL from the selected dilution was spread plated on the already set tryptic soy agar (TSA). Two (2) loopfuls of each of the bacteria from 3-day old cultures on TSA were each transferred separately to 50 mL tryptic soy broth (TSB) medium and incubated overnight at 28±2°C. A loopful from each TSB bacterial inoculum was streaked on prepared TSA to have pure isolates from each broth culture. Viability was confirmed by standard plate count method using tryptone soy broth plus 2% agar (TSBA). These inocula were prepared in order to use them *in vitro* for testing the antifungal and biocontrol activities of the isolates.

### 2.6 PGPR and biochemical analysis of bacteria isolates

#### 2.6.1 Detection of hydrogen cyanide (HCN) production

Bacterial isolates were screened for the production of hydrogen cyanide (HCN) production according to the methodology previously described by Castric [17]. Bacterial cultures were streaked on nutrient agar medium containing 4.4 g per liter of glycine. Picric acid solution (0.5% in 2% sodium carbonate) was prepared and Whatman filter paper No 1 was soaked in it and was placed inside the lid of a plate which was sealed with parafilm. After plates were incubated at 30°C for 4 days, production of HCN was observed by the light brown to dark brown colour that developed and no colouration development indicated negative activity.

#### 2.6.2 Determination of indole acetic acid (IAA) production

Fifty (50) mL of nutrient broth (Merck) containing 0.1% (D) L-tryptophan were inoculated with 500 μL of 24 h old bacterial cultures and incubated in refrigerated incubator Shaker at 30°C and 180 rpm for 48 h in the dark. The bacterial cultures were then centrifuged at 10,000 rpm for 10 min at 4°C [18].

An aliquot of 1 mL of supernatant was transferred into a fresh tube to which 50 μL of 10 mM orthophosphoric acid and a 2 mL of Salkowski reagent comprising (1 mL of 0.5 M FeCl3 in 50 mL of 35% HCIO4) were added. The mixture was incubated at room temperature for 25 min. The development of a pink colour indicated the presence of indole acetic acid [19]. The absorbance of the pink solution from each isolate was measured and recorded at 530 nm using spectrophotometer (Thermo Spectronic, Merck, SA).

#### 2.6.3 Determination of Phosphate solubilisation (PS)

Bacterial isolates were spot inoculated on Pikovskaya agar medium plates. The plates were incubation at 28°C for 7 days. Phosphate solubilization activity was observed as clear zone around the colonies while no zone was considered negative activity [20].

#### 2.6.4 Detection of ammonia (NH_3_) production

Peptone water was used to determine ammonia production of bacterial cultures. Freshly grown cultures were inoculated into 10 mL peptone water and incubated for 48–72 h at 30°C. Nessler’s reagent (0.5 mL) was added in each tube after incubation and positive test was observed as brown to yellow colour development while negative activity was observed with no colour development [21].

#### 2.6.5 Determination of 1-aminocyclopropane-1-carboxylate (ACC)

This procedure was carried out according to the protocol of Li et al., 2011. Ninhydrin reagents were prepared and five working concentrations of ACC were used which are 0.05, 0.15, 0.2, 0.3 and 0.5 mmol^- 1^ for colorimetric assay using the 96-wells PCR plates. Absorbance was read at 570 nm.

#### 2.6.6 Catalase activity

A sterile toothpick was used to mix 48 h old bacterial colonies placed on a clean glass slide to which a drop of 3% hydrogen peroxide was added. The effervescence that follows indicated catalase positive activity while no effervescence indicated negative activity.

#### 2.6.7 Assay for protease production

Extracellular protease production was assayed according to Maurhofer, Keel, Haas and Défago [22]. Spot inoculation of each bacterial isolate on skim milk agar plate was carried out and incubated at 37°C for 24 h. Development of halo zone around the bacterial colony was considered as a positive test for protease production while absence of halo zone was considered negative test.

#### 2.6.8 Oxidase activity

Oxidase activity was determined by using the filter paper spot method [23]. Kovács oxidase reagent (1-2 drops) was added to 24 h old culture on a small piece of filter paper. Change in colour to dark purple within 60 to 90 s was considered as oxidase positive test while absence of colour change indicated negative activity.

All analysis were repeated twice

### 2.7 Antifungal effect assay

Potato dextrose agar (PDA) medium was used as the medium to assay for the antifungal activities of 8 isolates against *F. graminearum* which is a toxin producing fungi and pathogenic to man, animals and plants. This was carried out by inoculating the pathogenic fungi at the centre of the medium and then streaking the isolates on the medium 3 cm away from the fungi. The clear zones between isolates and fungi after incubation for 4 to 7 days at room temperature indicated antagonist interaction between them.

#### 4.2.8 Antibacterial effect assay

Antagonistic activity of isolates against *B. cereus* and *E. feacalis* was carried out. *B. cereus* is a food poisoning pathogen worldwide with serious health implication, so it is very important in the food industry and agriculture [24]. *E. feacalis* is among the diverse type of bacteria capable of causing infection in man, animals and plants causing death of plant within 7 days [25]. They were screened by using a perpendicular streak method (Parthasarathi et al., 2010). In perpendicular streak method, Luria Bertani agar (Merck) was used and each plate was streaked with test bacterial isolates at the centre of the plate and incubated at 30°C for 48 h to allow optimum growth. Later, 24 h fresh sub-cultured isolated bacteria were prepared and streaked perpendicular to the test isolates and incubated at 37°C for 24 h. The experiment was carried out in triplicate.

### 2.9 Isolation of genomic DNA

Genomic DNA of all isolates was extracted using ZR soil Microbe DNA MiniPrep™ (Zymo Research, USA) extraction kit. Bacterial cultures were grown in 10 mL of Luria Bertani broth (Merck) at 37°C for 24 h and then centrifuged at 10,000 rpm (Universal Z300K model centrifuge; HERMLE Labortechnik, Germany) for 5 min. The bacterial pellets were resuspended in 200 μL of distilled water and transferred to ZR Bashing Bead™ lysis tube and 750 μL lysis solutions were added to the tube. The bashing bead was secured in a bead beater fitted with a 2-mL tube holder FastPrep^®^ 24 and processed at a maximum speed for 5 min. The ZR BashingBead™ lysis tube was centrifuged in a microcentrifuge at 10,000×g for 1 min, 400 μL of supernatant was transferred to a Zymo-Spin™ IV Spin Filter in a collection tube and was centrifuged at 7,000×g for 1 min and 1,200 μL of Soil DNA Binding Buffer was added to the filtrate in the collection tube. Eight hundred (800) micro litres of the mixture of the binding buffer and filtrate was transferred to a Zymo-Spin™ IIC Column in a collection tube and centrifuged at 10,000×g for 1 min. Two hundred (200) μL of DNA Pre-Wash Buffer was added to the Zymo-Spin™ IIC column in a new collection tube and centrifuged at 10,000×g for 1 min. Five hundred (500) μL soil DNA Wash Buffer was added to the Zymo-Spin™ IIC Column in a new collection tube and centrifuged at 10,000×g for 1 min. The Zymo-Spin™ IIC Column was transferred to a clean 1.5 mL micro-centrifuge tube and 100 μL DNA Elution Buffer was added directly to the column matrix. The tube was centrifuged at 10,000×g for 30 s to elute the DNA.

### 2.10 PCR Amplification targeting the 16S rRNA

PCR was carried out in 25 μL reaction volumes. Each reaction contained 12.5 μL of PCR master mix, 0.5 μL of primer, 11 μL of nuclease free water and 1.0 μL of DNA template. The 16S rDNA gene was amplified using the universal bacterial primers F1 (5′-GAGTTTGATCCTGGCTCAG-3′) and R2 (5′- GWATTACCGCGGCKGCTG-3′) (Maynard et al 2005). PCR was performed using a DNA Engine DYAD™ Peltier thermal cycler (Bio-Rad). The PCR program used was an initial denaturation at 95°C for 5 min, followed by 35 cycles of denaturation at 94°C for 30 s, annealing at 62°C for 30 s and extension at 72°C for 1 min, followed by a final extension at 72°C for 5 min.

### 2.11 Agarose gel electrophoresis procedure

One times Tris-Acetate EDTA (1X TAE) buffer was prepared by adding 4900 mL distilled water to 100 mL of 50X TAE (375 mL of Tris-Cl, 28.55 mL of acetic acid, 50 mL of EDTA and 46.45 mL distilled water) and filled in the electrophoresis tank. 1.5% agarose gel (4.5 g in 300 mL of 1X TAE buffer) was prepared by melting in microwave until boiling. Once the agarose was dissolved, it was allowed to cool to ~45°C before pouring into the casting mold. Before pouring the agarose gel, combs of desired sizes were inserted into the tray in such a way that no bubbles were caught under the teeth. After the gel had cooled, the combs were gently removed. The gel was placed in the electrophoresis tank to which 150 μL ethidium bromide has been added to 1.5 L buffer to cover it to a depth of about 1 cm. DNA samples were prepared by mixing 10 μL of the PCR reaction mixture with 10 μL of 1X loading buffer in 1:1 ratio on sterile parafilm. Twenty (20) μL samples were loaded into each well in the gel with a sterile micropipette and taking care not to cross-contaminate the wells. Six (6) μL of molecular marker (1 kb DNA ladder (Fermentas)) was loaded in the first and the last wells of each comb. The voltage was set to 80 V and top cover was attached, making sure that the polarity of the preparation was placed correctly. After about 1 h when the loading dye had migrated to the mid-point of the gel, the power was turned off and the gel removed. DNA fragments were visualized by removing the gel slab from the tray and placing it on a UV trans- illuminator. The outcome of running the gel was recorded/captured using Chemidoc^™^ MP imaging system (Bio-Rad USA).

### 2.12 Sequencing and phylogenetic analysis

The Sequencing of the purified PCR products was conducted at facilities of Inqaba Biotechnical Industrial (Pty) Ltd, Pretoria, South Africa using ABI PRISM^®^ 3500XL DNA Sequencer (Applied Biosystems). Only isolates with best activities in terms of biochemical, biofertilization and biocontrol were sequenced. The chromatograms were edited using Chromas Lite version 2.4 software (Technelysium Pty Ltd 2012) Nucleotide sequences were analyzed and edited by using BioEdit software (Hall, 1999). The obtained 16S rDNA sequences were compared to sequences in the NCBI GenBank database with the Basic Alignment Search Tool (BLAST) (Altschul et al., 1990). Multiple alignments of the sequences were carried out by Mafft programme 7.050 (Katoh, 2013) against corresponding nucleotide sequences of the genus selected isolate retrieved from GenBank. Phylogenetic analyses were conducted using software’s in MEGA version 5.2.2 (Tamura et al., 2011). Evolutionary distance matrices were generated as described by (Jukes and Cantor, 1969) and a phylogenetic tree was inferred by the neighbor-joining method (Saitou and Nei, 1987). Tree topologies were evaluated by bootstrap analysis (Felsenstein, 1985) based on 1000 resamplings of the neighbor-joining data set. Manipulation and tree editing were carried out using TreeView (Page, 1996).

### 2.13 Supporting data

The 16S rDNA gene sequences determined for the bacterial isolates in this study were deposited into the GenBank database and assigned accession numbers (Table 4).

## 3.0 Results

### 3.1 Physical and chemical characterization of samples

#### 3.1.1 Cation Exchange Capacity (CEC), Nitrate and Nitrogen analysis of soil samples

In Figure 2, it was observed that the exchangeable cations and the nitrate value followed the same trend in the graph. They decreased from the 4 WAP to the 8 WAP and increased again at the 12 WAP which was the peak and decreased again at the 16 WAP only to start increasing gradually to the time of harvest. The nitrate content ranged between 6.28 mg/kg at the 8 WAP and 26.16 mg/kg at the 4 WAP. From the 12 WAP, it decreased again from 19.06 mg/kg to 6.40 mg/kg at the 16 WAP and gradually increased again to the time of harvest to 7.39 mg/kg.

**Figure 2:**
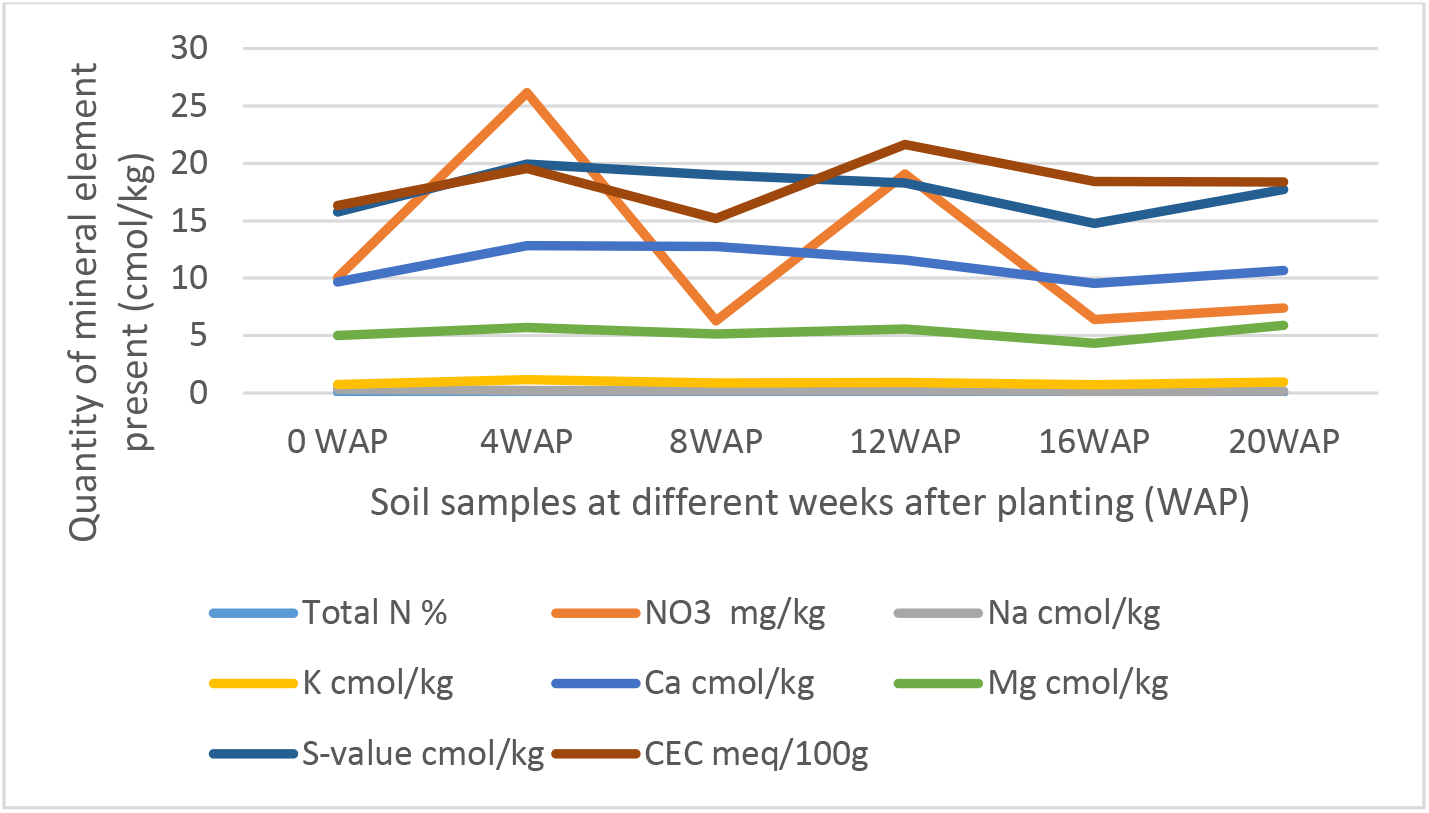
Comparison of different physical and chemical properties of soil samples between 4 and 20 weeks after planting (WAP). N=nitrogen, K=potassium, Ca=calcium, Na=sodium, Mg=magnesium, S-value shows the mean of the whole graph

All chemical parameters, N, K, Ca, Na and Mg increased from zero the original bulk soil to 4 WAP. Cation Exchangeable capacity (CEC) and Nitrate followed the same pattern of decreasing from the time of fertilization/flowering (4 WAP) till the 8 WAP and increasing at 12 WAP. They gradually decreased from 12 WAP to 16 WAP and gradually increased again to the time of harvest. Samples at 12 WAP had the highest CEC of 21.64 meq/100g. The lowest was 15.19 meq/100g at 8 WAP. Nitrate (NO_3_) was highest at 4 WAP and lowest at 8 WAP. After 4 WAP, all the minerals kept decreasing gradually until the 16 WAP (which is the time of maturity of seeds) and then started increasing gradually again. The only exceptions are the individual cations Mg, Na, K and Ca which all followed the same pattern of gradually decreasing up until the 16^th^ week and then gradually increasing afterwards to the time of harvest (Fig. 3 and Fig. 4).

**Figure 3:**
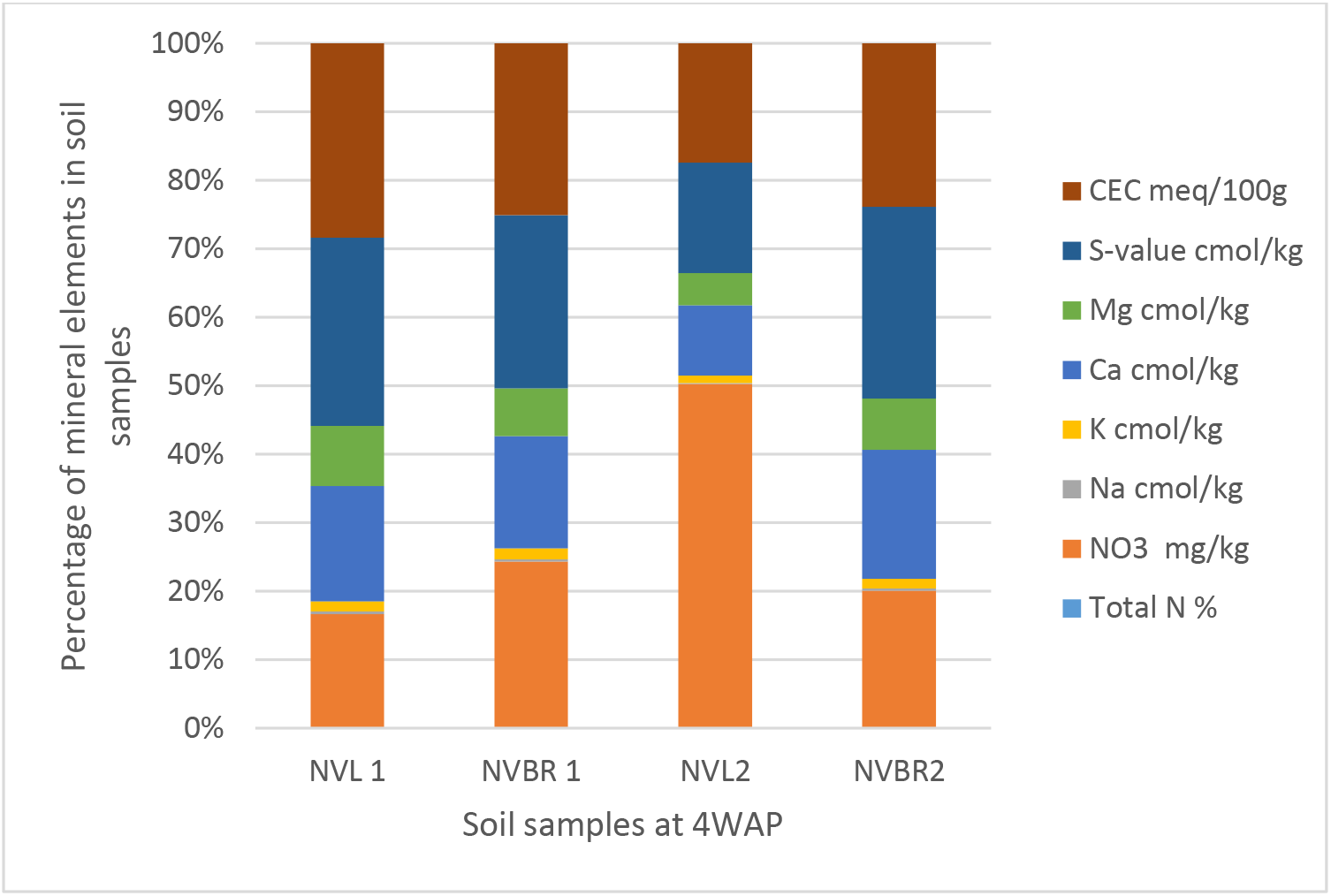
Physical and chemical analysis of soil samples at 4 WAP. Nitrate (NO3) and CEC were the highest quantity of mineral found in the soil samples and both were found in NVL2 and NVL1 respectively. N=November; VL and VBR=Variety of Bambara from whose root, soil samples (NVL1, NVL2, NVBR1, NVBR2) were taken

**Figure 4:**
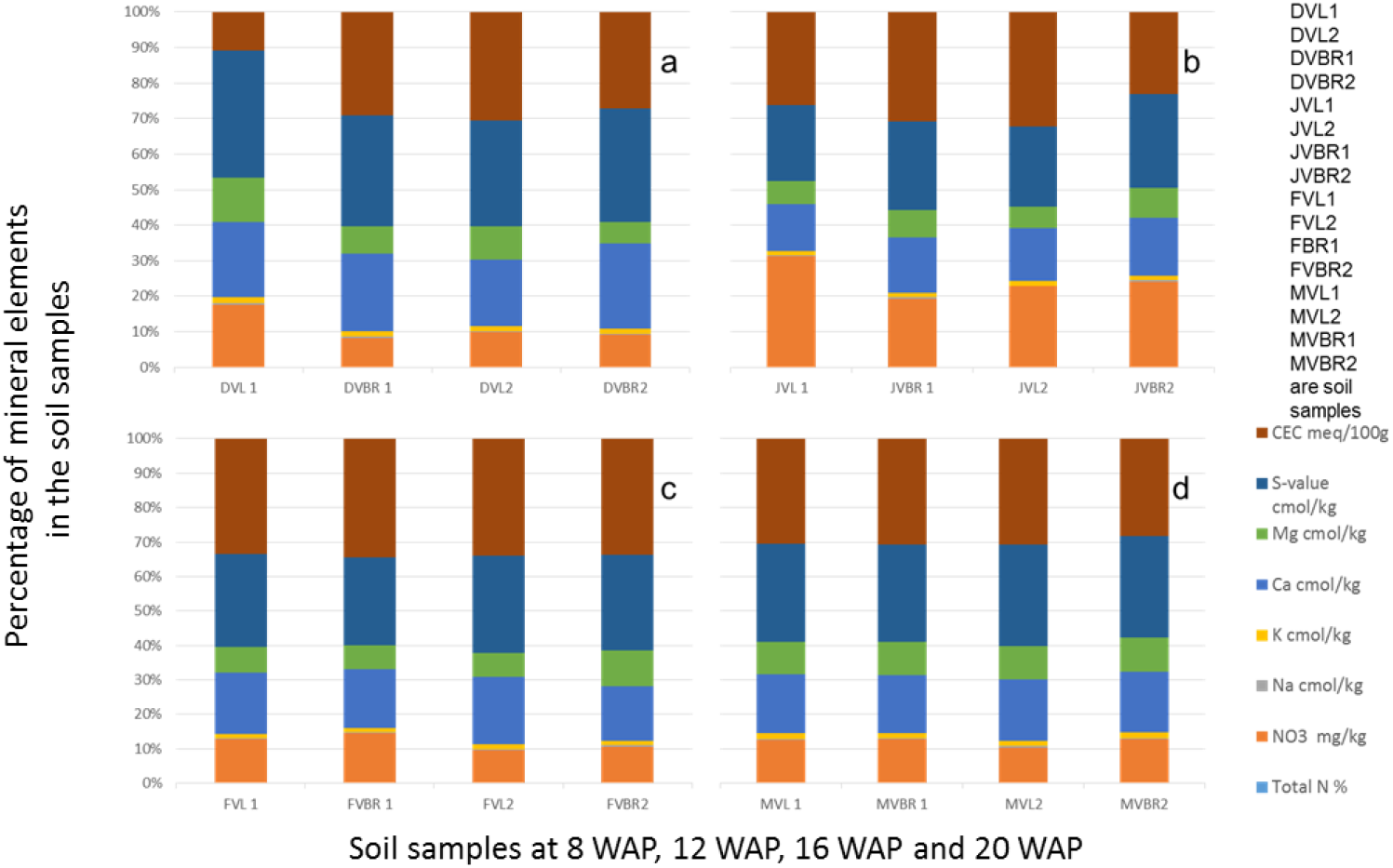
Physical and chemical analysis of soil samples at 8, 12, 16 and 20 WAP. DVL1, DVL2, DVBR1, DVBR2 are soil samples from 8 WAP (a), JVL1, JVL2, JVBR1, JVBR2 are soil samples from 12 WAP (b), FVL1, FVL2, FVBR1, FVBR2 are soil samples from 16 WAP (c), MVL1, MVL2, MVBR1, MVBR2 are soil samples from 20 WAP (d), D=December; J=January; F=February; M=March; VL and VBR=Variety of Bambara from whose root soil samples were taken

Furthermore, Figure 4 (a) revealed that CEC and Ca were the highest mineral found in the soil samples and both were found in DVL2 and DVBR2 while NO3 had decreased and total N is seen to increase from the previous weeks where it was absent in all samples. (b) showed that CEC and NO_3_ were the highest mineral found in the soil samples and both were found in JVL1 and JVL2 (c) showed that CEC is highest and found in FVBR1 while NO_3_ has decreased again but available in all samples. (d) showed that CEC is highest and found in MVBR1 while NO_3_ has gradually increased again and available in all samples. Also total N increased and consistent in b, c, d samples.

#### 3.1.2 pH and Redox tolerance of soil samples

The different soils at the different growth stages differed in their physical and chemical properties. The pH reduced gradually from the original bulk soil to the soil at the time of harvest. The pH ranged from 2.3 at 16 WAP which was the lowest, to 3.4 at 12 WAP which was the highest. The reduction-oxidation relationship of both living and non-living things is measured by the redox potential (Eh) measured in volts. The Eh in this study ranged from 170.33 mV at 12 WAP to 221.67 mV at 16 WAP. This pattern of Eh and pH shows that they are both negatively correlated. This result reveals that nitrogen fixation by root nodule bacteria and activities of rhizospheric bacteria can affect the condition of the soil from one season to the other (Fig. 5a and Fig. 5b). Also though redox value, <300 mV can be limiting for plant growth but bambara groundnut has grown and also increased in yield.

**Figure 5:**
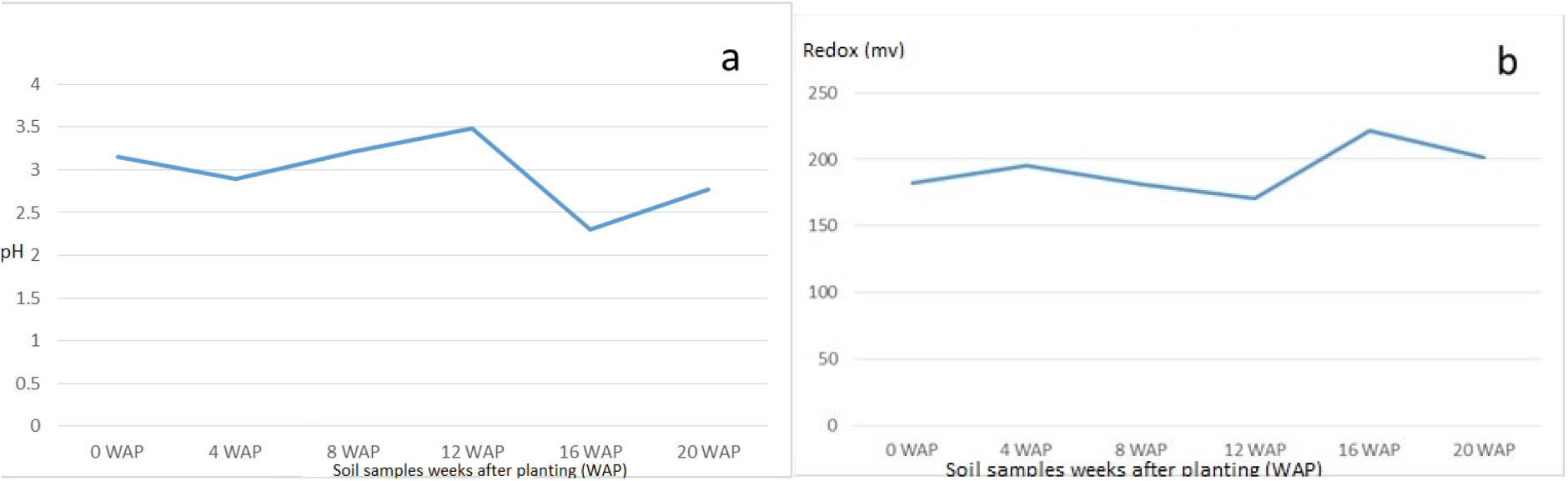
**(a)** Average pH values of soil samples from original soil to the time harvest. The highest pH was at 12 WAP while the least was at 16 WAP. **(b)** Average Redox value for soil samples at the different growth stages from original soil to the time harvest. The highest value was at 16 WAP while the lowest was at 12 WAP. The line graph for redox and pH are inversely related.

### 3.2 Organic matter content of soil samples

The organic matter of soils varies based on the type of soil. The organic matter in this study ranged from 2.02% at 0 WAP (bulk soil) which is the lowest to 3.46% which is the highest at 16 WAP. The organic matter kept increasing form 4 WAP to 16 WAP and reduced at the time of harvest. Total organic carbon also followed the same pattern as the total organic matter. It ranged from 1.18% at 0 WAP which was the lowest to 20.01% at 16 WAP which was the highest (Figure 6).

**Figure 6:**
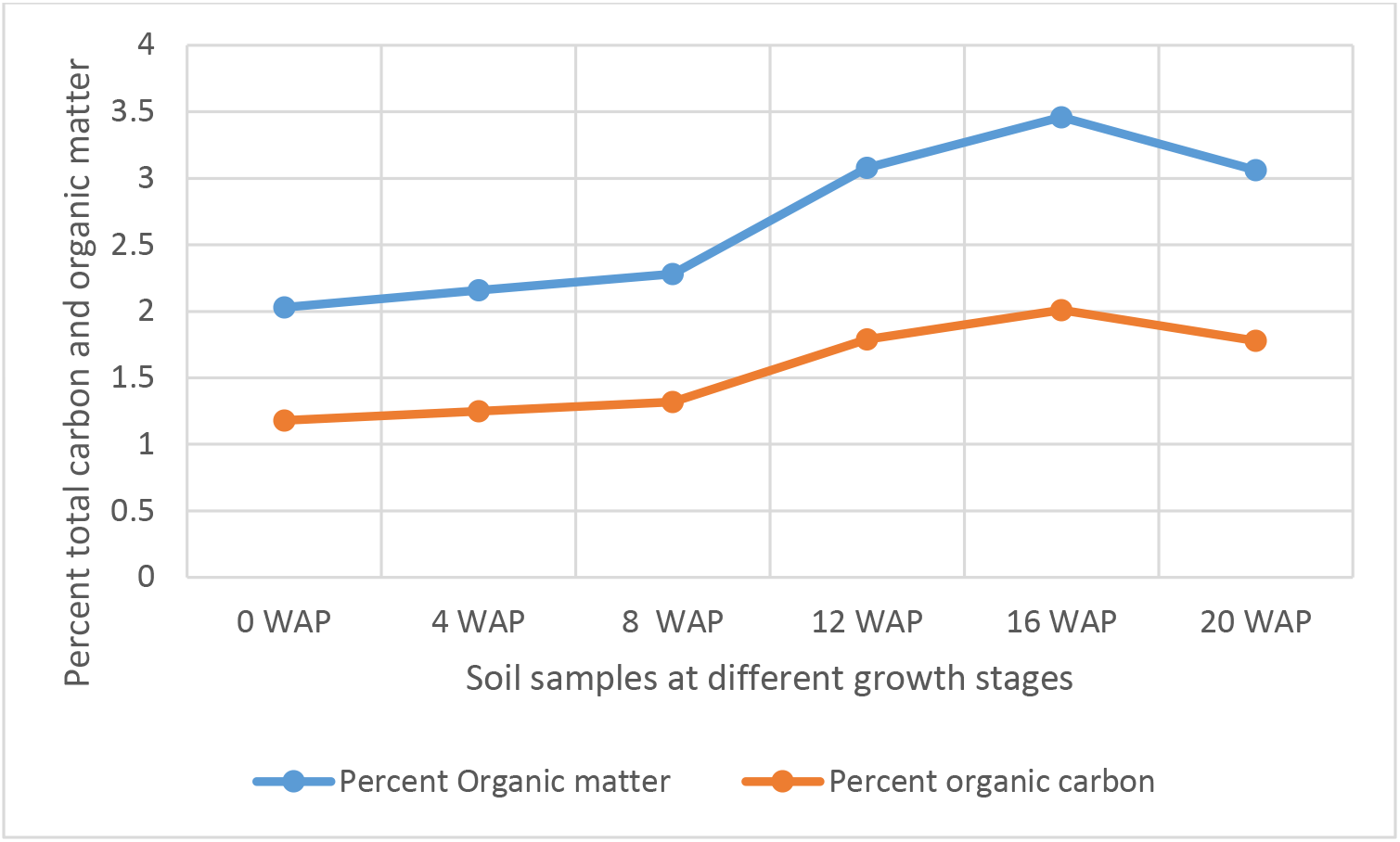
Carbon and organic matter content of soil samples from 0 WAP to harvest (20 WAP)

### 3.3 Culturing and isolation of bacteria from soil samples

The 43 isolates subcultured spanned through the different growing seasons. Some isolates that grew at the beginning of the growth period were also isolated at harvest while most of them were not. Most of the organisms isolated at the time of harvest were not the same as those that were there from beginning. The number of isolates increased from 4 WAP to 8 WAP only to decrease at 12 WAP and then continued increasing up till the time of harvest. The period of the 12 WAP corresponded to the pod and seed formation period (Table 1).

**Table 1:** Total number of different isolates by morphology at different growth stages

| Soil Samples |  | Soil Samples |  | Soil Samples |  | Soil Samples (16 |  | Soil Samples (20 |  |
| --- | --- | --- | --- | --- | --- | --- | --- | --- | --- |
| (4 WAP) | *NOI | (8 WAP) | *NOI | (12 WAP) | *NOI | WAP) | *NOI | WAP/harvest) | *NOI |
| NVL 1 | 3 | DVL 1 | 5 | JVL 1 | 3 | FVL 1 | 3 | MVL 1 | 22 |
| NVBR 1 | 2 | DVBR 1 | 9 | JVBR 1 | 2 | FVBR 1 | 4 | MVBR 1 | 17 |
| NVL2 | 8 | DVL2 | 8 | JVL2 | 5 | FVL2 | 11 | MVL2 | 8 |
| NVBR2 | - | DVBR2 | 2 | JVBR2 | 3 | FVBR2 | 4 | MVBR2 | - |
| **Total** | **13** |  | **24** |  | **13** |  | **22** |  | **47** |

\*NOI is number of isolates

### 3.4 PGPR activities

From the PGPR tests, 41.87% (18) showed positive actions in two or more of the PGP tests. Out of which 4.65% of the isolates were positive for HCN production, all were positive for NH_3_ production and ACC with absorbance at 570 nm and standard curve was drawn (Fig 7a and Fig 7b). Isolates that were positive for IAA production were 16.28%. Absorbance value was recorded for all organisms (Fig 8a) and 27.91% solubilized phosphate (Table 2). Standard curve for IAA was also plotted (Fig 8b). This was used to calculate the quantity of IAA produced by each isolate (Appendix). Of all the isolates, 27.91% showed positive activity for catalase, oxidase and protease production but the isolates that were positive in at least two of the PGP tests were all positive in at least one of the biochemical test while 13.95% of the isolates that were not positive to at least 2 of the PGP tests were positive to all the 3 biochemical tests (Table 3). While out of the 18 isolates used in this study, 27.77% were positive to catalase, oxidase and protease production.

**Figure 7a:**
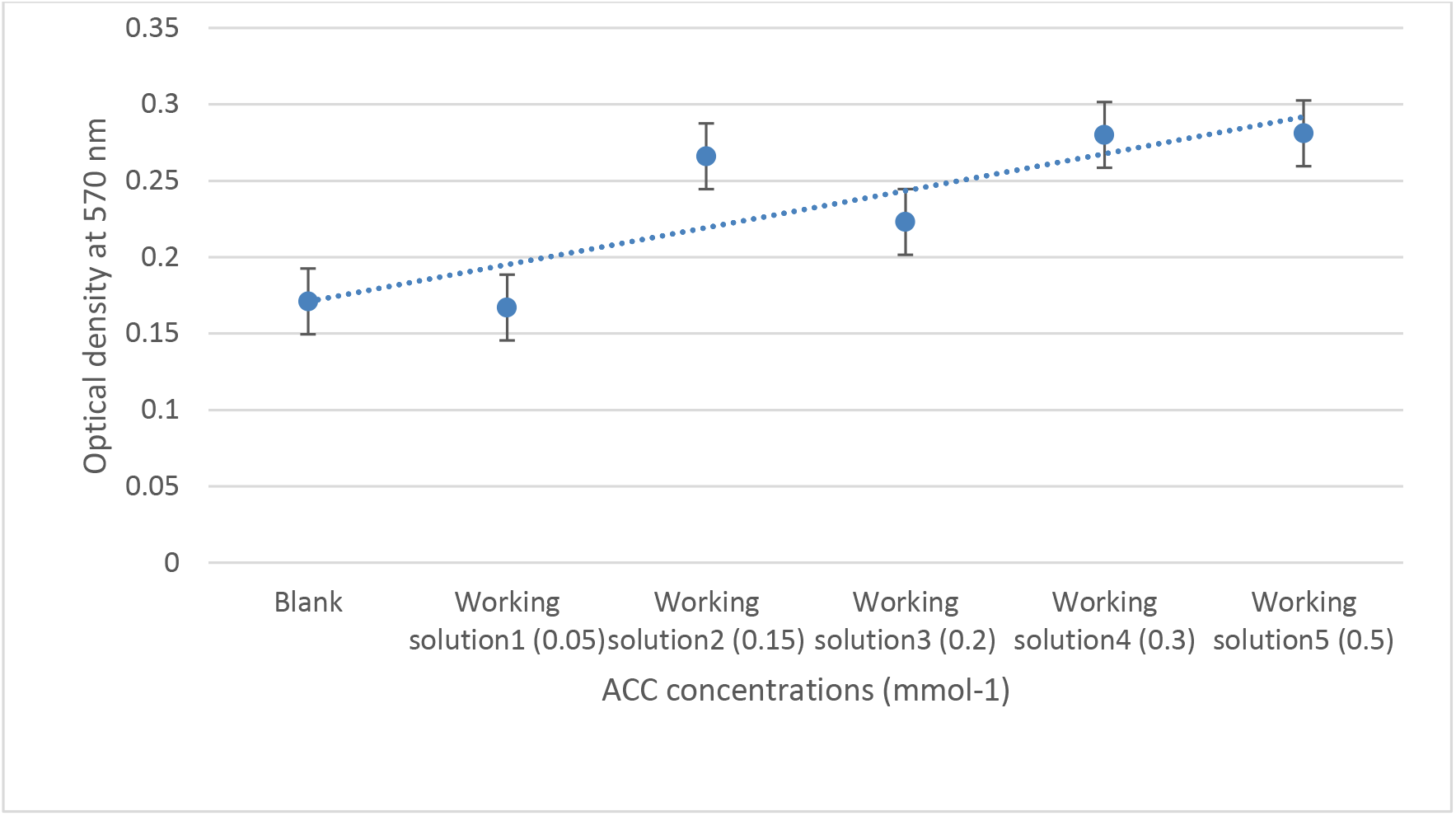
Standard curve of ACC concentrations ranging from 0.05 to 0.5 mmol^-1^determined by the 96- well PCR-plate ninhydrin assay (y=0.0242x+0.1467, R^2^ = 0.7364). Each data point represents the mean from triplicate determinations, and the error bar represents the standard error ACC, 1-aminocyclopropane- 1-carboxylate.

**Figure 7b:**
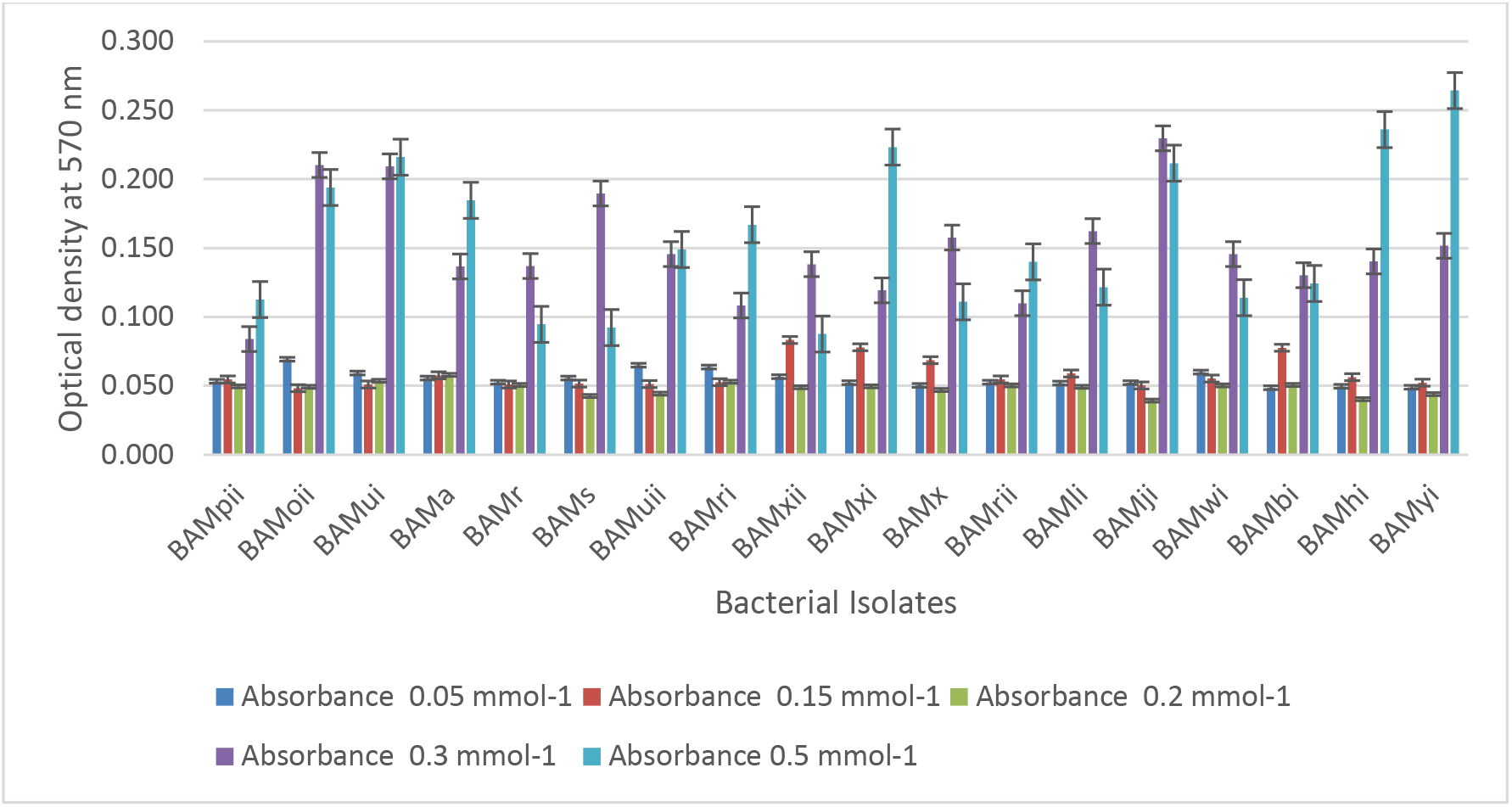
Absorbance of bacterial isolates at different concentration of ACC

**Figure 8a:**
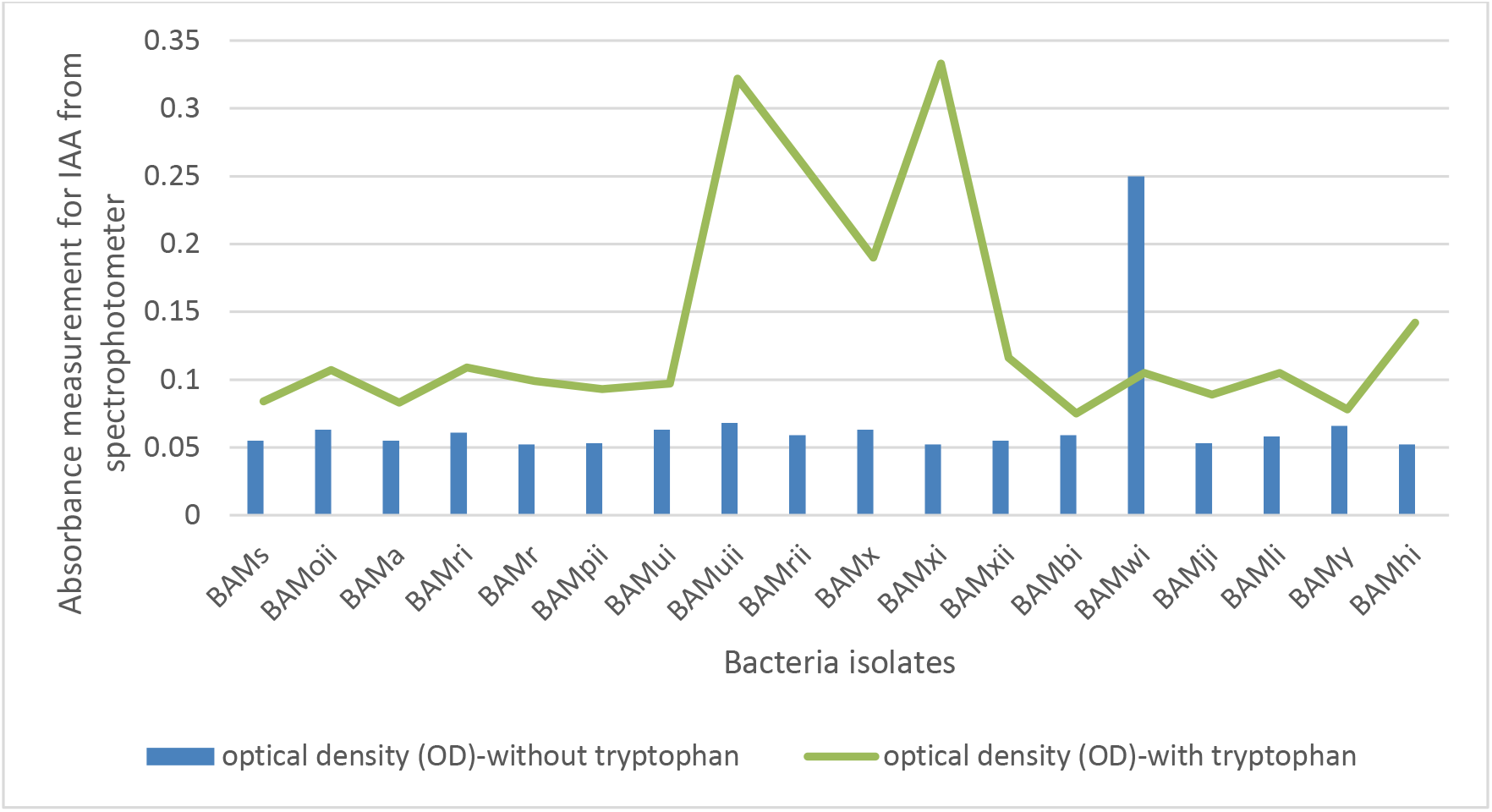
Spectrophotometric measurement of absorbance of IAA in Isolates in the presence or absence of tryptophan at optical density of 530nm

**Figure 8b:**
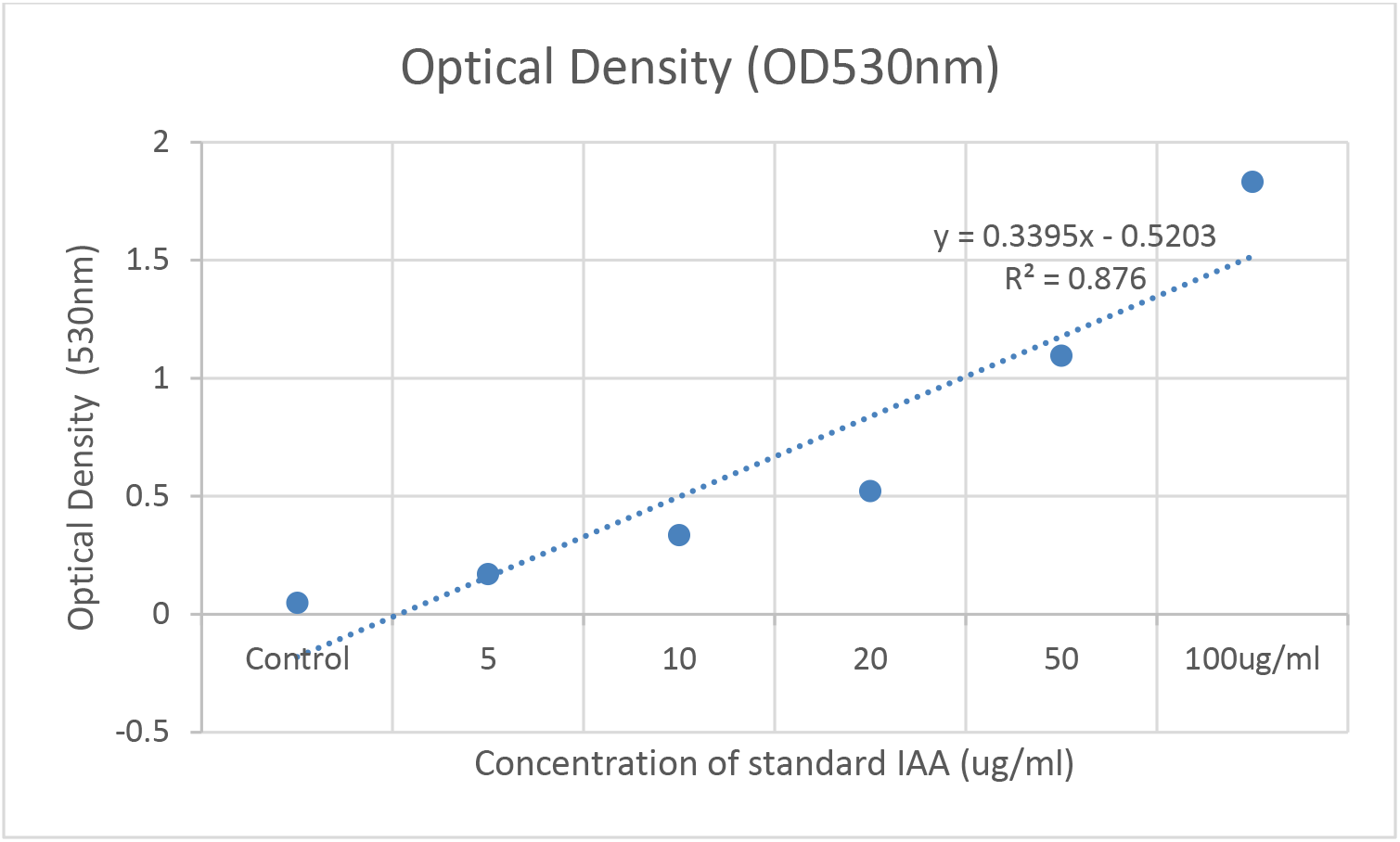
Standard graph of IAA at optical density of 530nm

**Table 2:**
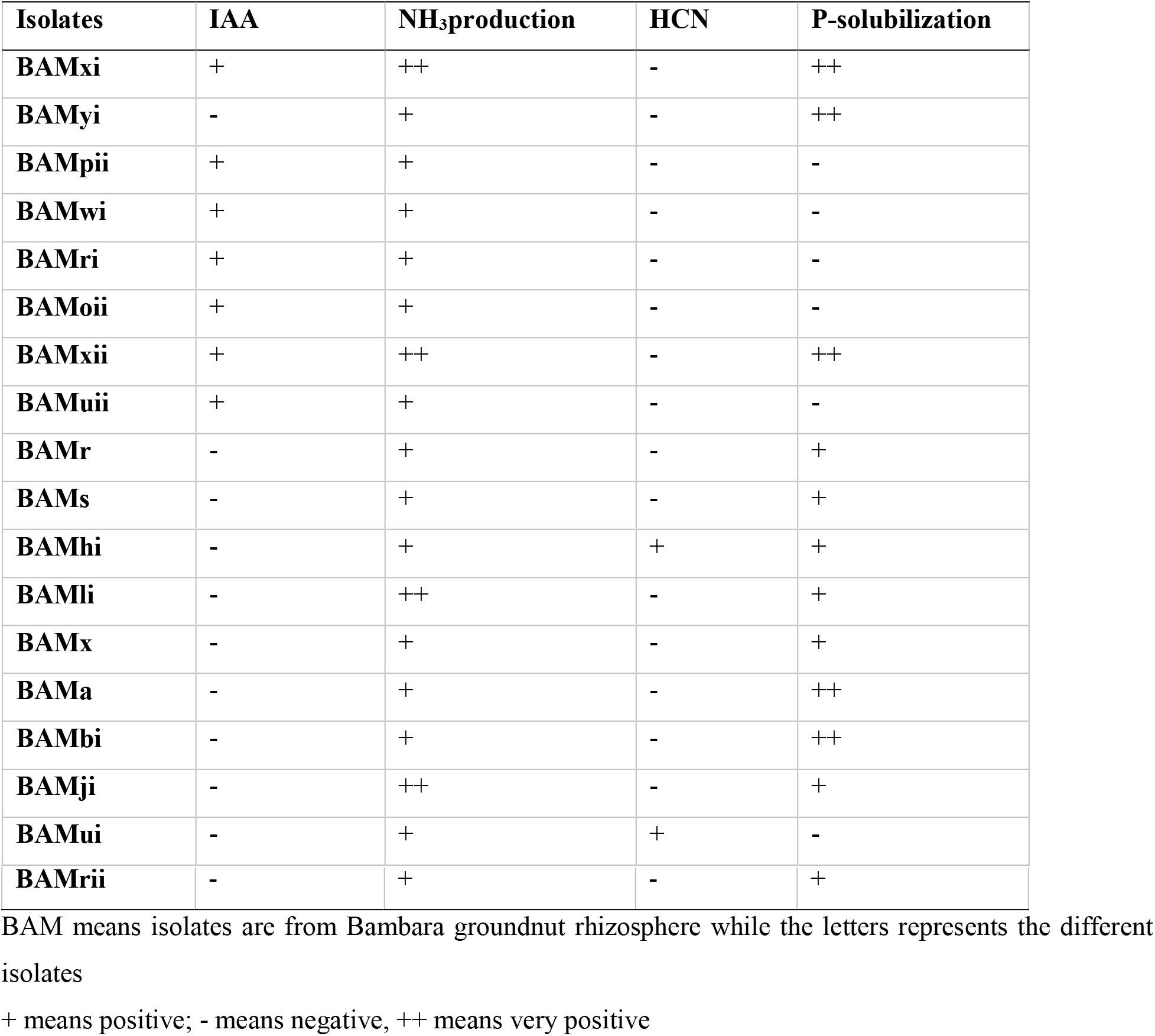
Plant growth-promoting activities of rhizobacteria isolates from the above-mentioned soil samples

**Table 3:** Biochemical activities of bacterial isolates from Bambara groundnut rhizosphere

| **Isolates** | **Catalase** | **Oxidase** | **Protease** |
| --- | --- | --- | --- |
| **BAMxi** | + | + | - |
| **BAMyi** | ++ | - | - |
| **BAMpii** | + | + | - |
| **BAMwi** | + | + | - |
| **BAMri** | + | - | - |
| **BAMoii** | + | + | - |
| **BAMxii** | + | + | - |
| **BAMuii** | ++ | - | - |
| **BAMr** | ++ | + | + |
| **BAMs** | ++ | + | + |
| **BAMhi** | + | - | - |
| **BAMli** | ++ | + | + |
| **BAMx** | - | + | + |
| **BAMa** | + | - | - |
| **BAMbi** | + | + | + |
| **BAMji** | + | + | + |
| **BAMui** | + | - | - |
| **BAMrii** | + | - | - |

BAM means isolates are from bambara groundnut rhizosphere while the letters represents the different isolates
+ means positive; - means negative, ++ means very positive

### 3.5 Biocontrol activities

Biocontrol activities were carried out in this study to test for the antifungal and antibacterial potentials of the bacterial isolates.

#### 3.5.1 Antifungal activities

Isolated bacteria were tested against *Fusarium graminearum* (written as f.g on the plate). BAMji, BAMr, BAMli and BAMhi (9.3%) (*B. cereus*, *B. amyloliquefaciens*, *B. thuringiensis, Bacillus sp.)* showed antifungal potential against *F. graminearum* (Fig. 9).

**Figure 9:**
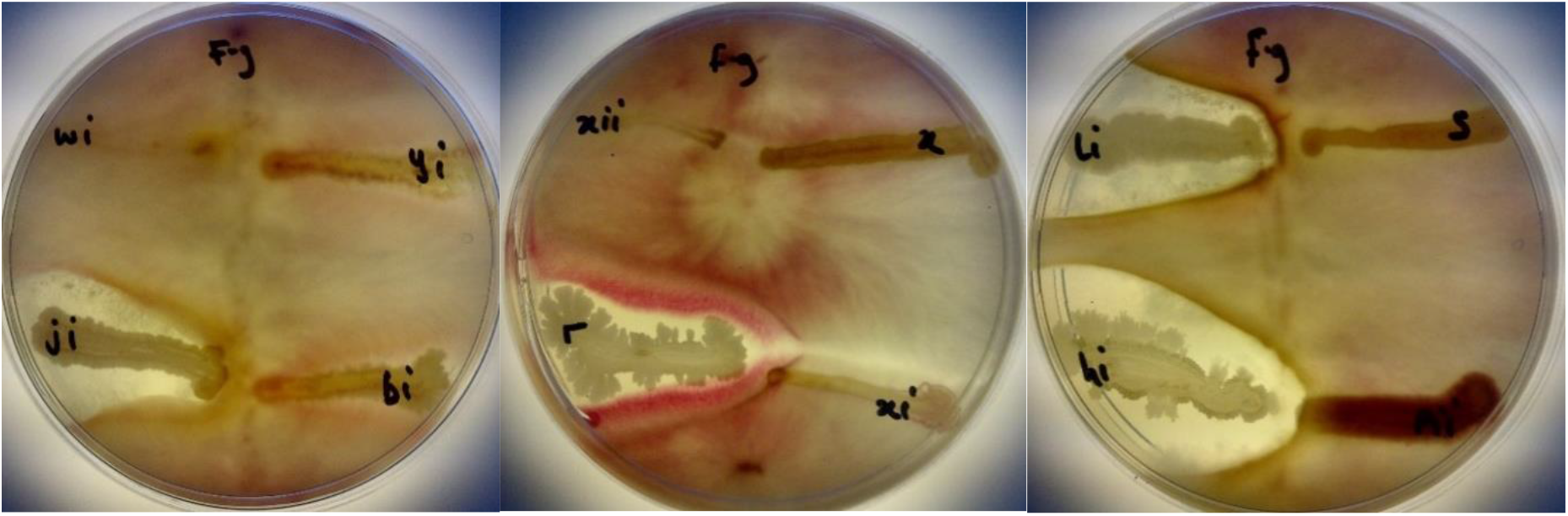
Antifungal activities of BAMji, BAMr, BAMli and BAMhi against *F. graminearum (Source: Ajilogba and Babalola 2019)*

#### 3.5.2 Antibacterial activities

Rhizobacteria from this study were tested against *B. cereus* (written as B.C on the plates) and *E. faecalis* (*written as* E.F *on the plates*). BAMui, BAMli, BAMoii, BAMyi, BAMhi, and BAMpii (16.2%) had antagonistic effects against *B. cereus* and *E. faecalis* as seen in the pattern formed on the streaked pathogen (Fig. 10).

### 3.6 Molecular identification of selected isolates

Bacterial universal primers (F1R2) amplified 1.5 kb fragment from the genomic DNA of the isolates. Computational analysis was used as a means of identifying the isolates. Analysis of the partial sequences of the 16S rDNA gene of the selected isolates was used as a means of identifying them at the genus level. The BLAST tool was used to compare the partial nucleotide sequences of the 16S rDNA gene of the isolates with the nucleotide database of NCBI web server. From the BLAST search, it was inferred that the isolates were members of the GC-rich firmicutes, actinobacteria and proteobacteria.

The 16S rDNA gene sequence of the selected isolates was obtained by BLASTn search; however, 27 strains of combination of the phylum firmicutes, actinobacteria and proteobacteria were selected based on high identity (%) with good E value. Table 4 results show that query sequences were best pairwise aligned with 16S rDNA gene sequence of other firmicutes, proteobacteria and actinobacteria with sequence similarity and identity ranged between 96-99%, with E-value of 0.

**Figure 10:**
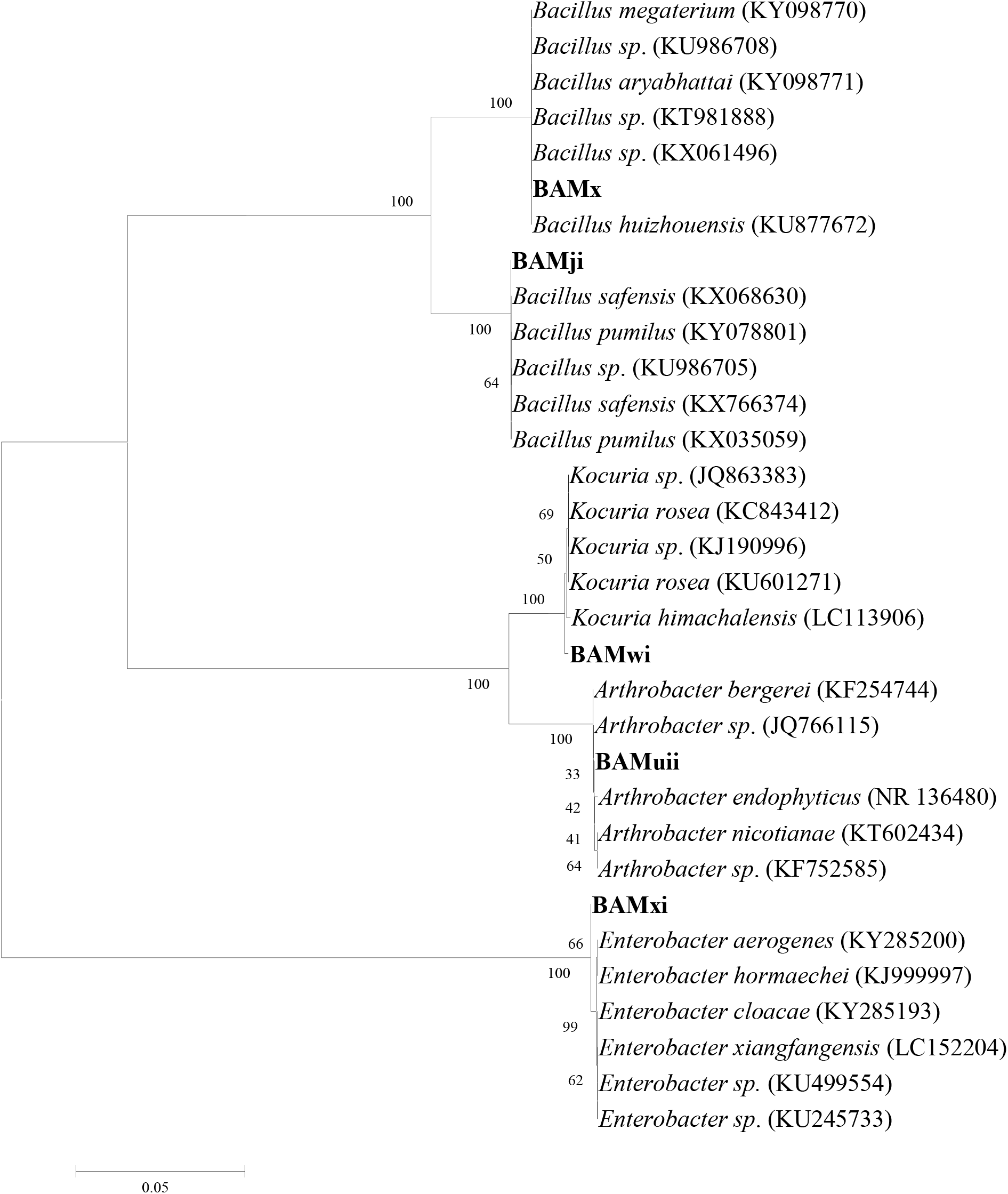
Phylogenetic tree based on 16S rRNA sequences using neighbour-joining method for bacterial isolates and their closely related type strains.

**Table 4:** Results of 16S rDNA gene sequence similarities of rhizobacteria isolates and GenBank accession numbers using BLASTn algorithm isolate code

| Isolate code | Sequence length (bp) | Closest related strain in database | Accession number | Similarity (%) | E-value |
| --- | --- | --- | --- | --- | --- |
| BAMji | 1424 | *B. cereus* | KX588094 | 99 | 0 |
| BAMr | 1058 | *Bacillus sp.* | KX588095 | 98 | 0 |
| BAMui | 1405 | *Bacillus sp.* | KX588096 | 97 | 0 |
| BAMx | 1417 | *B. megaterium* | KX588099 | 99 | 0 |
| BAMuii | 1353 | *Glutamicibacter (Arthrobacter) bergerei* | KX588097 | 98 | 0 |
| BAMwi | 1335 | *Kocuria rosea* | KX588098 | 99 | 0 |
| BAMxi | 1377 | *Enterobacter hormaechei* | KX588100 | 99 | 0 |
| BAMa | 1387 | *Staphylococcus sp.* | KX588093 | 96 | 0 |

### 3.7 Phylogenetic analysis and diversity

The isolates were subjected to phylogenetic analysis. The 16S rDNA sequences of the bacterial isolates were aligned with reference nucleotide sequences obtained from the GenBank. The phylogenetic position of the bacterial isolates was evaluated by constructing a phylogenetic tree using neighbour joining method (Fig. 11). This method placed the bacterial isolates in different clades encompassing members of their genera; this was supported with bootstrap values. Bootstrap values based on 1000 replications were listed as percentages at the branching points.

**Figure 11.**
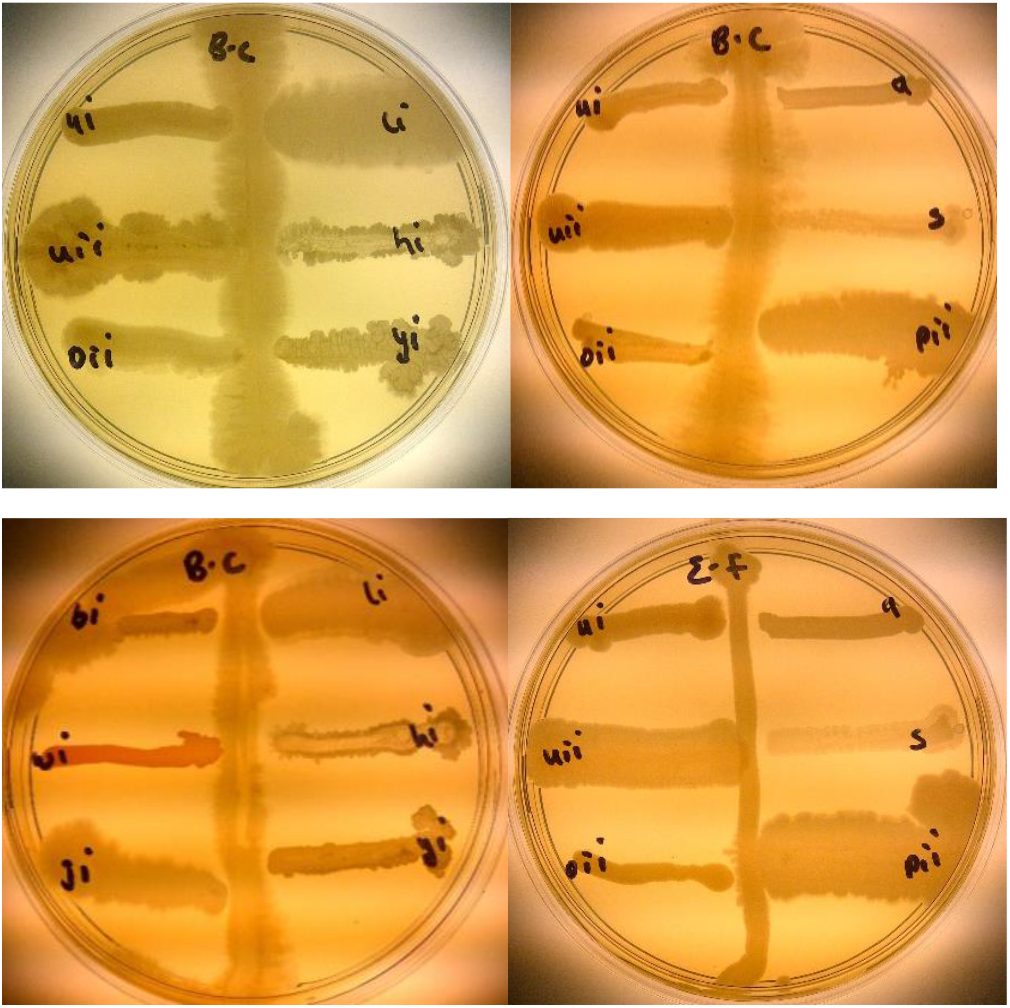
Antibacterial activities of isolates BAMui, BAMli, BAMoii, BAMyi, BAMhi, and BAMpii against *B. cereus* and *E. faecalis*

## 4.0 Discussion

The plants’ rhizosphere, especially those of legumes, have been observed to be a microbiome for diverse PGPR. Bambara groundnut as a legminous plant is not only good for food but its rhizosphere is also very important in promoting plant growth and increasing food production. The rhizosphere is rich in total nitrogen, nitrate and CEC. In this study, the highest nitrate and CEC values were 26.16 cmol/kg at 4 WAP and 21.64 meq/100g at 12 WAP respectively while the least values were 6.28 cmol/kg and 15.19 meq/100g both at 8 WAP respectively (Fig. 4.3). The nitrate available in the soil falls in the same range with the soil nitrate analysed at different growing season from Dobra Voda and Chvalina farms during Spring and Autumn ranging between 4 and 14 ppm N, though some of the values from this study are higher [26]. These higher values of nitrate can be as a result of bambara groundnut’s ability to fix nitrogen through symbiosis with the bacteria in the root nodules. According to Vaněk, Šilha and Němeček [26], it can also be as result of the activities of leguminous crops in which case bambara groundnut is one. The CEC of soil samples in this study (between 15.19 and 21.64 meq/100g) fall within the range of CEC of organically and conventionally managed apple orchard ranging from 19.23 to 20.28 cmol/kg [27], even though no fertilizer or organic manure was applied to bambara groundnut in this study.

The variation in the physical and chemical composition of the soil at different growth stages of bambara groundnut shows the involvement of these rhizobacteria in nitrification (Fig. 3, Fig. 4 and Fig. 5). This is because as the number of rhizobacteria increases (Table 1), the cmol/kg of nitrate decreases, this could be as a result of uptake of nitrate by plants through the rhizobacteria [28]. As the number of rhizobacteria increases, the usage of NO_3_ increases by the plant and so its availability in the soil decreases. This is in agreement with the observation made that with an increase of PGPR in the soil, plant uptake of nitrogen from fertlizer applied to the soil also increases [29]. This is obvious in 8 WAP and 16 WAP where the number of isolates were high but the volume of NO_3_ and other CEC were low. It could also mean that though the rhizobacteria were available in the soil they might not be nitrogen fixers. Nitrogen available in the soil can also be low if plant density is low since both are positively correlated and might not be as a result of poor symbiotic relationship [30]. This was contrary to other findings which observed that nitrogen increased with increase in microbial population and activities [31]. In the 12 WAP, when NO_3_ was very high but number of isolates was low, it corresponded to the period of pod and seed formation. This helps to shed more light on the reason for the increase in NO_3_ and also if the type of rhizobacterial available were nitrogen fixers or the number of nitrogen fixers increased in the soil at this growth stage. Apart from nitrogen-fixers being in the nodules there are other free living rhizobacteria that are non-symbionts, free living in the rhizosphere and can also fix nitrogen [32] even though in small amounts [33].

Normally, leguminous plants are sensitive to pH and this is important to their nitrogen-fixing ability because at low pH the soil is acidic and inhibits the activities of nitrogen-fixing bacteria and nitrogen is not released for plant growth [34]. Bambara groundnut grows well on acidic soils [35] which agrees with the result from this study. It was observed that there was an increase in alfalfa yield when there was no additional nitrogen and the pH was high which is in contrast to the observation in this study. Although the pH of the soils throughout the season were acidic as they ranged from 2.3 to 3.49, nitrogen was available in the form of nitrate as mentioned earlier and bambara groundnut was enhanced in growth. Out of 12 isolates from the root nodule of bambara groundnut isolated from Cameroonian soil, 8 of them were able to grow at pH 3.5 in the soil which was acidic. This also reveals the potential of bambara groundnut to grow under very harsh condition [36]. Soil pH and redox potential (Eh) are negatively correlated, as one increases the other decreases. The Eh in this study ranged from 170.33 to 221.66 mV which falls into the category of moderately reduced soils having Eh between +100 and +400 mV and close to cultivated soils with Eh range of between +300 and +500 mV [37]. The Eh and pH of the soil help to determine the type of metabolism evident in the bacterial community of the soil and invariably the biological activities of the soil. Growth and development of soil bacteria and their metabolic, enzymatic and microbial activities are directly or indirectly affected by the Eh and/or the pH [38, 39].

Soil organic matter in the soil is very vital and important as it can produce as much carbon found in the combination of that of the atmosphere as carbon dioxide and the biomass of plants [40]. In this study, the soil organic matter increased till 16 WAP and decreased at harvest, this could be as a result of heavy metabolic processes involving pod and seed formation at the 12 and 16 WAP. Assertion from this study points to the fact that between 4 WAP and 16 WAP, the soil of Bambara groundnut kept increasing and at 16 WAP experienced the highest level of fertility. At this point, other cereals can be cropped with Bambara groundnut to improve their growth.

Bacteria from the rhizosphere are important in auxin production and vitamin synthesis that encourage biofertilization [41]. Bacterial isolates from this study were able to produce IAA and solubilise phosphate which are important in biofertilization to increase crop growth. Ammonia, HCN and siderophore production and phosphate solubilization are also able to contribute to biocontrol potentials of rhizobacteria [21]. Seventy eight percent (78%) of the isolates from different monocotyledonic plants’ rhizosphere and soil [21], 77.1 % of isolates from chickpea rhizosphere [42], 85.57% of isolates from maize rhizosphere [43] produced ammonia as against 100% ammonia production from this study. This could be as a result of the high nitogen-fixing capacity of both the nitrogen fixers and free living non-symbionts in the rhizosphere of bambara groundnut.

Soil phosphorous is an important macronutrient needed for plants to grow, hence deficiency of Phosphorus in soil is a major challenge in agricultural food/crop production. Total soil P occurs in either organic or inorganic form. The major form of organic phosphorus in the soil is phytate (salts of phytic acid). As a source of phosphorus, it constitutes about 60% of soil organicphosphorus and this organic form is poorly utilized by plants [44]. It either forms a complex with cations or adsorbs to various soil components, so it is not readily available to plants. For phytate-P to be immobilized so that it is easily utilized by plants, phytate has to be solubilized. Phytate can then be dephosphorylated by phosphatases (phytases and phosphatases) before it is assimilated by plants [45]. Some soil microorganisms secrete phytases into the soil to be able to make use of organic substances from plants roots and also when they compete with plant roots for elements such as phosphorus [46]. The phytases secreted into the rhizosphere invariably help to immobilize phytate and make it soluble so that it is utilised by the plants [47].

Phosphate solubilization is a complex phenomenon and it helps to discriminatively screen the bacteria which are able to break down tricalcium phosphate (TCP) and thereby release inorganic phosphate. According to Laslo, György, Mara, Tamás, Ábrahám and Lányi [21], 63.8% of bacteria isolates from different rhizosphere of monocotyledonous plants solubilized phosphate as against 27.91% from this study. It was observed that none of the isolates from fields growing chickpea from West of Allahabad Agricultural Institute, India produced HCN [42] while in this study 4.65% produced HCN. This result is not comparable to a report by Agbodjato, Noumavo, Baba-Moussa, Salami, Sina, Sèzan, Bankolé, Adjanohoun and Baba-Moussa [43] which revealed that 100% of isolates from maize rhizosphere produced HCN.

Isolates *B. cereus*, *B. amyloliquefaciens*, *B. thuringiensis, Bacillus sp.* from this study were able to antagonise the growth of *F. graminaerum,* which is an agricultural challenge to barley, wheat and maize in South Africa [48, 49]. It was observed that *B. amyloliquefaciens* produced HCN in addition to ammonia and solubilizing phosphate while *B. cereus*, *B. thuringiensis* and *Bacillus sp.* produced ammonia and solubilised phosphate. Isolates BAMui, BAMoii, BAMyi, BAMpii, *B. amyloliquefaciens* and *B. thuringiensis.* were able to antagonise the growth of *B. cereus* and *E. faecalis*. Three of the antifungal isolates also displayed antibacterial activities which show that some rhizobacteria are both antifungal and antibacterial agents while the rest are just purely antibacterial. *Lysobacter spp* strains have been found to carry out both antimicrobial and antifungal activities against *Pythium ultimum, Colletotrichum gloeosporioides, F. oxysporum, Botrytis cinerea, Rhizoctonia solani, Botryosphaeria dothidea and B. subtilis* [30]. The ability of these isolates to be able to inhibit and/or suppress the growth of both fungi and bacteria implies the richness of the bambara groundnut rhizosphere and its ability to resist diseases and pests.

It is important to identify bacterial isolates at the species level as vital information concerning the organisms such as its novelty and its ability to produce bioactive compound is provided (Adegboye and Babalola, 2012). The 16S rDNA gene sequence analysis was used to identify selected bacterial isolates in this study. The comparison of the bacterial isolates sequences revealed 96-99% identification similarities with 16S rDNA gene sequence of the genus *Bacillus, Enterobacter, Arthrobacter and Kocuria.* The 16S rDNA gene sequences analysis has been shown to be a very effective tool in phylogenetic characterization of microorganisms (Thenmozhi and Kannabiran, 2010). This analysis is important as it helps to explain the evolutionary relationship that exist among microorganisms. The phylogenetic relationship of the potent bacterial isolates to known *Bacillus, Arthrobacter, Enterobacter, and Kocuria* spp was first estimated through a BLAST search of the GenBank database. In order to have analysis that is robust and reliable, the strains that are closest to the selected isolates were chosen and used for comparison of pairwise sequence and also for the phylogenetic tree construction. The selected bacterial isolates were grouped distinctly in different branches. Sometimes strains of bacteria that are grouped distinctly produce distinct microbial agents (Intra et al., 2011).

The analysis of 16S rRNA nucleotide sequences is used to determine higher taxonomic relationships of microorganisms [50]. The nucleotide sequences comparison of the bacterial isolates showed 99-100% identification similarities with those of reference nucleotide sequences from the GenBank. The phylogenetic tool is a powerful method that helps to elucidate the evolutionary relationship among organisms [51]. The tree revealed that the bacterial isolates have a phylogeny that is well supported and completely resolved. It also shows high resolution of all inner branches. Overall, this phylogenetic tree with its high-level branching is in consonance with traditional systematic divisions. This is because organisms that belong to the same family or genus taxonomically are grouped into different species using the traditional systematic divisions.

### Relationship between Bambara groundnut physical and chemical analyses, pH, Eh, PGPR and biocontrol

Bambara groundnut physical and chemical analyses in this study reveal that the soil is an acidic soil and the growth of Bambara groundnut in the soil makes it more acidic and that it can thrive in acidic soil. It has been observed that redox reaction is very important to the biocontrol activity of plant pathogen by rhizobacteria [37]. This, they do by generating reactive oxygen species (ROS) in the plant and this serves as an antagonistic response of the plant to the pathogen and indirectly stresses the pathogen [52]. Hydrogen peroxide and other signals like salicyclic and glutathione have been observed to increase the resistance of plants to pathogens [53]. The isolates in this study were also able to release hydrogen peroxide in the catalase reaction, which might have also enhanced their ability to resist pathogen growth.

## 5.0 Conclusion

The rhizosphere of Bambara groundnut is very rich in terms of biotic and abiotic components. It is quite interesting that most studies on Bambara groundnut have been on its food production but not much in-depth study has gone into its rhizosphere which is able to enhance its food production potential and then take it to the next level. This study revealed that the physical and chemical properties of soil at different growth stages are different and they affected the number, types and diversities of bacteria of Bambara groundnut rhizosphere. The Eh and pH of the soil were very important in the diversity of the bacterial isolates from the rhizosphere. They affected the type and abundance of the bacterial isolates at each growth stage. They were also important in the biocontrol potential of bacterial isolates. It is also observed from this study that PGP activities of rhizobacteria from Bambara groundnut’s rhizosphere is comparable to those of other legumes and crops. Also, that Bambara groundnut has great potentials in food security as biofertilizer and biocontrol agent against fungal and bacterial pathogens. These bacteria would be explored for their VOCs and how they can be used in other biotechnological processes.

## Acknowledgement

We gratefully acknowledge the North-West University for bursary to the first author and the National Research Foundation, South Africa, for grant that supports work in our laboratory.

